## Supplementary Information for "Comparative metabolomics with Metaboseek reveals functions of a conserved fat metabolism pathway in *C. elegans*"

Maximilian J. Helf[1,#], Bennett W. Fox[1,#], Alexander B. Artyukhin[2], Ying K. Zhang[1], Frank C. Schroeder[1,*]

Addresses

[1]Boyce Thompson Institute and Department of Chemistry and Chemical Biology, Cornell University, Ithaca, New York 14853, United States

[2]Chemistry Department, College of Environmental Science and Forestry, State University of New York, Syracuse, New York 13210, United States

[#]These authors contributed equally to this work

### Table of Contents

This file includes Supplementary Figures 1-13, Supplementary Tables S1-S4, and the NMR Appendix.

Supplementary Tables S5 and S6 available as a separate file.

Supplementary Figures........................................................................................................S3
Supplementary Tables........................................................................................................S17
NMR Appendix........................................................................................................S20

### Supplementary Figures

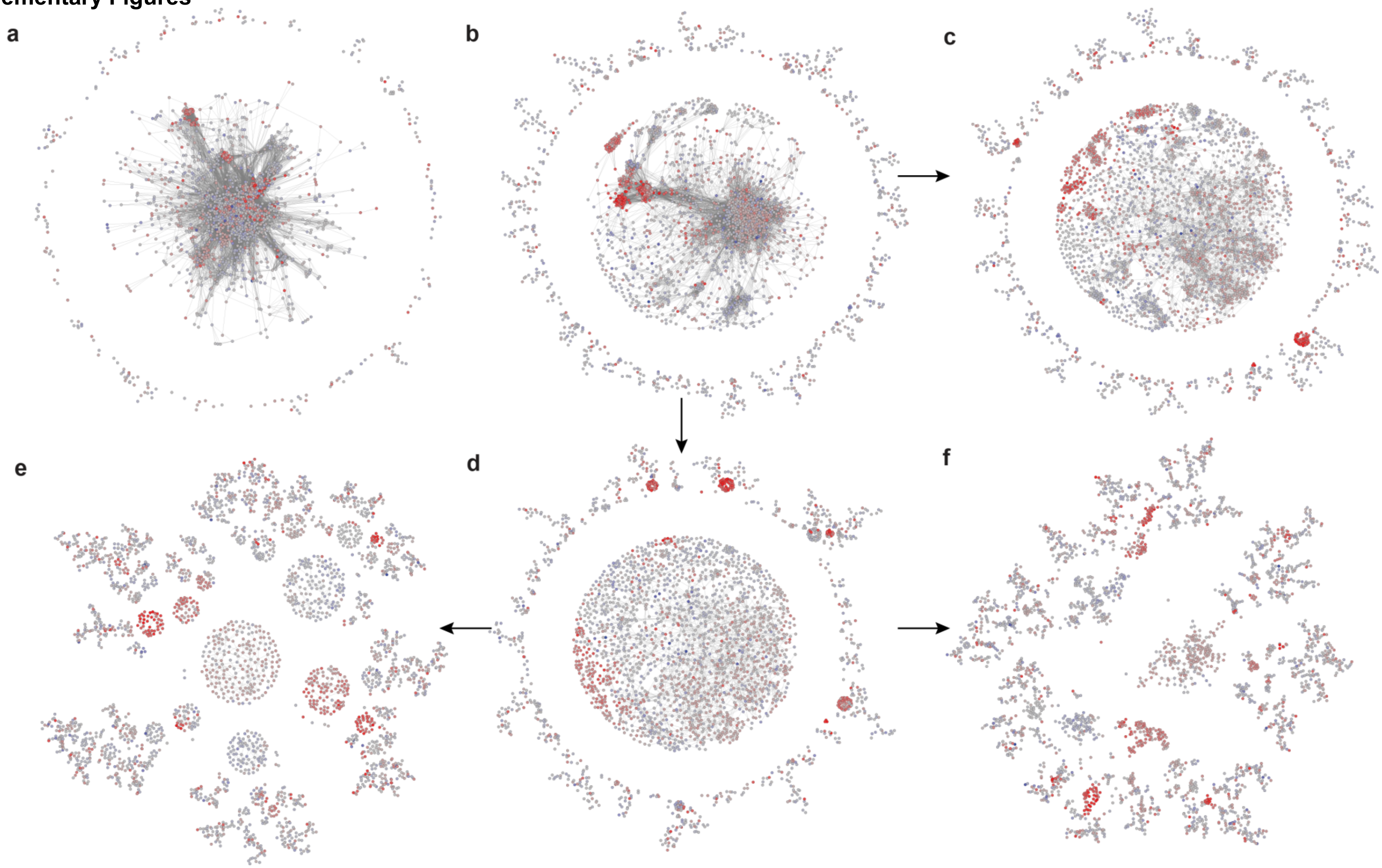

**Supplementary Figure 1**. Modifying MS/MS networks with the *Simplify Network* function. **a,** Negative ion MS/MS network constructed using a similarity score threshold of 0.6 and requiring a minimum of two matching fragments. **b,** Negative ion MS/MS network constructed using a similarity score threshold of 0.6 and requiring a minimum of four matching fragments. **c,** Simplification of the network in (**b**) by restricting the number of edges per node to the top 20, ranked by similarity score. **d,** Simplification of the network in (**b**) by restricting the number of edges per node to the top 10, ranked by similarity score. **e,** Further simplification of the network in (**d**) by restricting the maximum cluster size to 200 nodes, using the Metaboseek default display setting. This MS/MS network was rearranged slightly to save space and is featured in the main text (see **Figure 4**). **f,** Simplification of the network as in (**e**), but using the Metaboseek *KK* display setting.

**a**

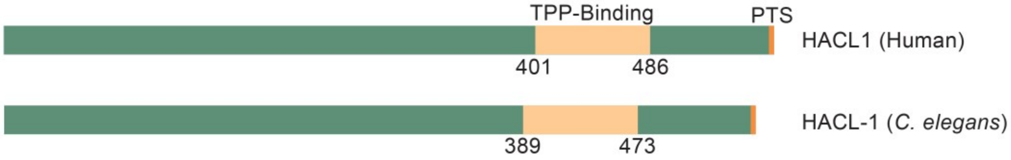

**b**

```
Human       MPDSNFAERSEEQVSGAKVIAQALKTQDVEYIFGIVGIPVTEIAIAAQQLGIKYIGMRNE
C. elegans  -------------MDGASITAAALKSQGVEYMFGVVGFPVIEVGMAAQAHGIKYIGCRNE
                         :.**.: * ***:*.***:**:**:** *:.:***  ****** ***

Human       QAACYAASAIGYLTSRPGVCLVVSGPGLIHALGGMANANMNCWPLLVIGGSSERNQETMG
C. elegans  QAAAYAAQAMGYLTGKPVALLVVSGPGILHAIGGLANATVNCWPVVCIGGTADVDLENRG
            ***.***.*:****.:* . *******::**:**:***.:****:: ***::: : *. *

Human       AFQEFPQVEACRLYTKFSARPSSIEAIPFVIEKAVRSSIYGRPGACYVDIPADFVNLQVN
C. elegans  AFQEWSQQESVRNSCKHVSRPTSLHTIPAHIEKAVRCAMYGRPGAVYVDLPGNLVLTSTE
            ****: * *: *   *. :**:*:.:**  ******.::****** ***:*.::*  ..:

Human       VNSIKYMERCMSPPISMAETSAVCTAASVIRNAKQPLLIIGKGAAYAHAE-ESIKKLVEQ
C. elegans  EEISFPPQVPLPAPVSIPPIAEIEKAIDTLKSAKKPLVIVGKGAAWSERGATQVQQFLTK
             :     :  :  *:*:   : : .* ..::.**:**:*:*****::.    .::::: :

Human       YKLPFLPTPMGKGVVPDNHPYCVGAARSRALQFADVIVLFGARLNWILHFGLPPRYQPDV
C. elegans  SKLPWLATPGGKGVASDLHPRFIGQARSLALREADTVFLIGARFNWILHFGLPPRFQKDV
             ***:* ** ****. * **  :* *** **: **.:.*:***:***********:* **

Human       KFIQVDICAEELGNNVKPAVTLLGNIHAVTKQLLEELDKTPWQYPPESKWWKTLREKMKS
C. elegans  KVVQIDICPEEFHQNVKTEVPLLGDIGETLAELTPRL--GDWTYDESTEWFKKLRENAEK
            *.:*:*** **: :***  * ***:*  .  :*  .*    * *  .::*:*.***: :.

Human       NEAASKELASKKSLPMNYYTVFYHVQEQLP-RDCFVVSEGANTMDIGRTVLQNYLPRHRL
C. elegans  NRAAVEKFVDDHSTPLNYYAAYQPIREFLANNDVIVINEGANTMDIGRTMMPSRLPKRRL
            *.** ::...:* *:***:.:  ::* *  .* :*:.***********:: . **::**

Human       DAGTFGTMGVGLGFAIAAAVVAKDRSPGQWIICVEGDSAFGFSGMEVETICRYNLPIILL
C. elegans  DAGTFGTMGVGHGFSLAAALWARDHSPKTKVLVVQGDSAFGFSAMELETIARYNLPVVTV
            *********** **::***: *:*:**   :: *:********.**:***.*****:: :

Human       VVNNNGIYQGFDTDTWKEMLKFQDATAVVPPMCLLPNSHYEQVMTAFGGKGYFVQTPEEL
C. elegans  IINNSGIYRGLLPEDDKAI--EGDRTLALPVLSLTAECRYEEMCKAFGGAGTVVRTVPEI
            ::**.***:*:  :  * :    * * .:* :.*  :.:**:: .**** * .*:*  *:

Human       QKSLRQSLADTTKPSLINIMIEPQATRKAQDFHWLTRSNM 578
C. elegans  KAALEKAFQKTDGPTVINALISTDSERKPQADHWLTRSKM 563
            : :*.::: .*  *::** :*. :: ** *  ******:*
```

**Supplementary Figure 2.** *C. elegans hacl-1 (*formerly *B0334.3*) encodes a protein similar to human HACL1. **a,** Protein structure schematic of human HACL1 and *C. elegans* HACL-1. Both proteins contain C-terminal TPP-binding domains and a peroxisomal targeting signal. **b,** Basic local alignment search tool (BLAST) comparison of human HACL1 and *C. elegans* HACL-1. Proteins are 49% identical overall, 64% identical in the annotated TPP-binding domain (highlighted) and share a type 1 peroxisomal targeting signal (boxed). Asterisks (*) indicate identical, two dots (:) indicates very similar, and one dot (.) indicates somewhat similar amino acid composition.

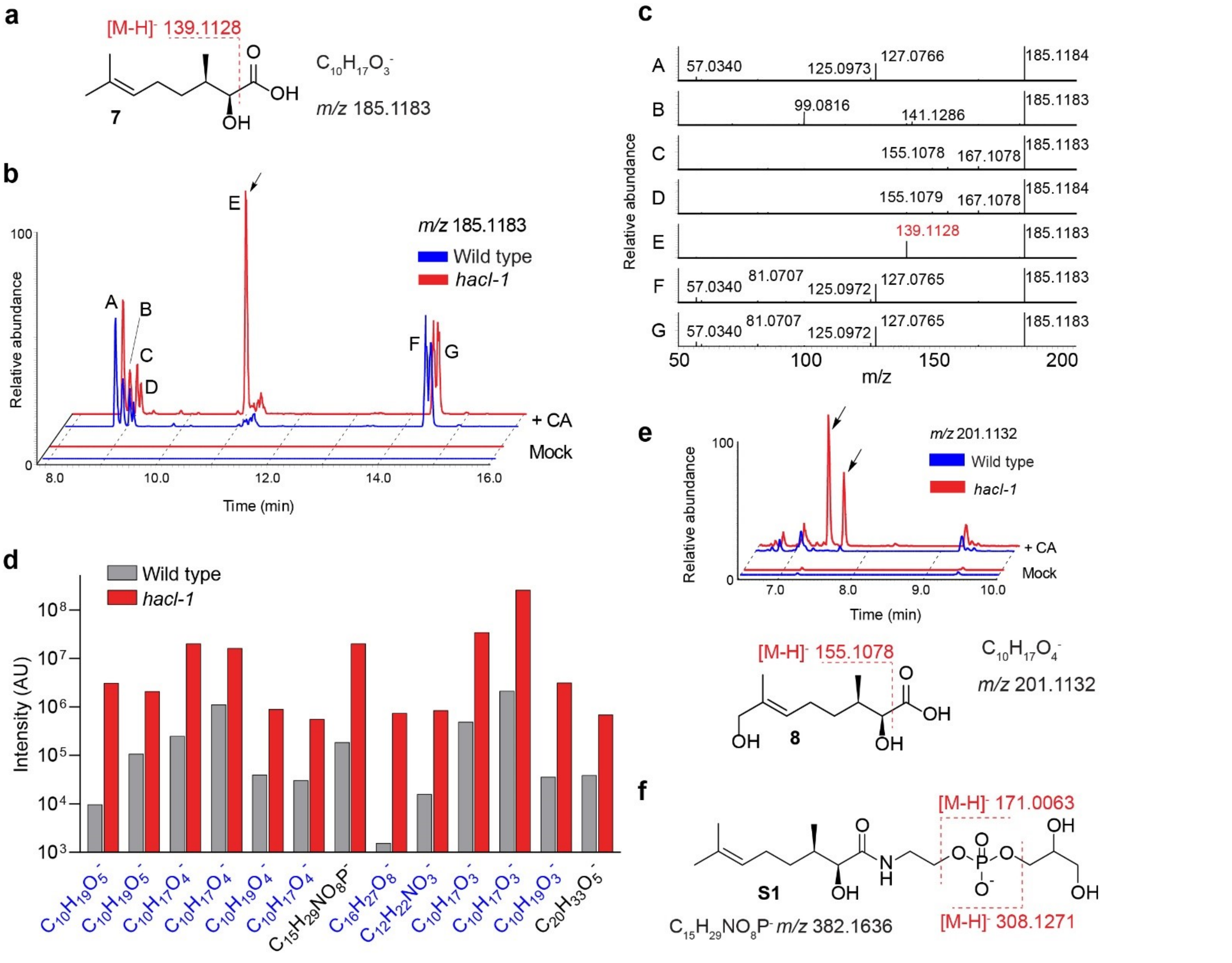

**Supplementary Figure 3.** Accumulation of α-hydroxy citronellic acid derivatives in *hacl-1*. **a**, Proposed structure of shunt metabolite **7** predicted to accumulate in *hacl-1* following CA supplement. **b**, EIC for *m/z* 185.1183 ($C_{10}H_{17}O_3^-$) reveals seven distinct CA-dependent metabolites (labeled A-G), but only feature E is enriched in *hacl-1* as compared to WT. **c**, MS/MS spectra for metabolites A-G. MS/MS fragmentation of E in negative ion mode produces a strong product ion with *m/z* 139.1128, corresponding to neutral loss of formic acid. **d**, Quantification of CA-derived metabolites in *hacl-1* that are at least $5x10^5$ intensity and 10-fold enriched in *hacl-1* as compared to wildtype animals, organized by increasing RT. Data represent one experiment. Metabolites in blue exhibit fragmentation between the carbonyl- and α-carbon in MS/MS. **e**, EIC for *m/z* 201.113 ($C_{10}H_{17}O_3^-$) reveals several distinct CA-dependent metabolites, two of which are enriched in *hacl-1* (marked with arrows). Both of these metabolites lose formic acid by MS/MS, proposed representative structure **8** below. **f**, Proposed structure and major MS/MS fragmentation reactions for CA-derived *N*-acyl glycerophosphoethanolamide (**S1**) accumulating in *hacl-1* larvae.

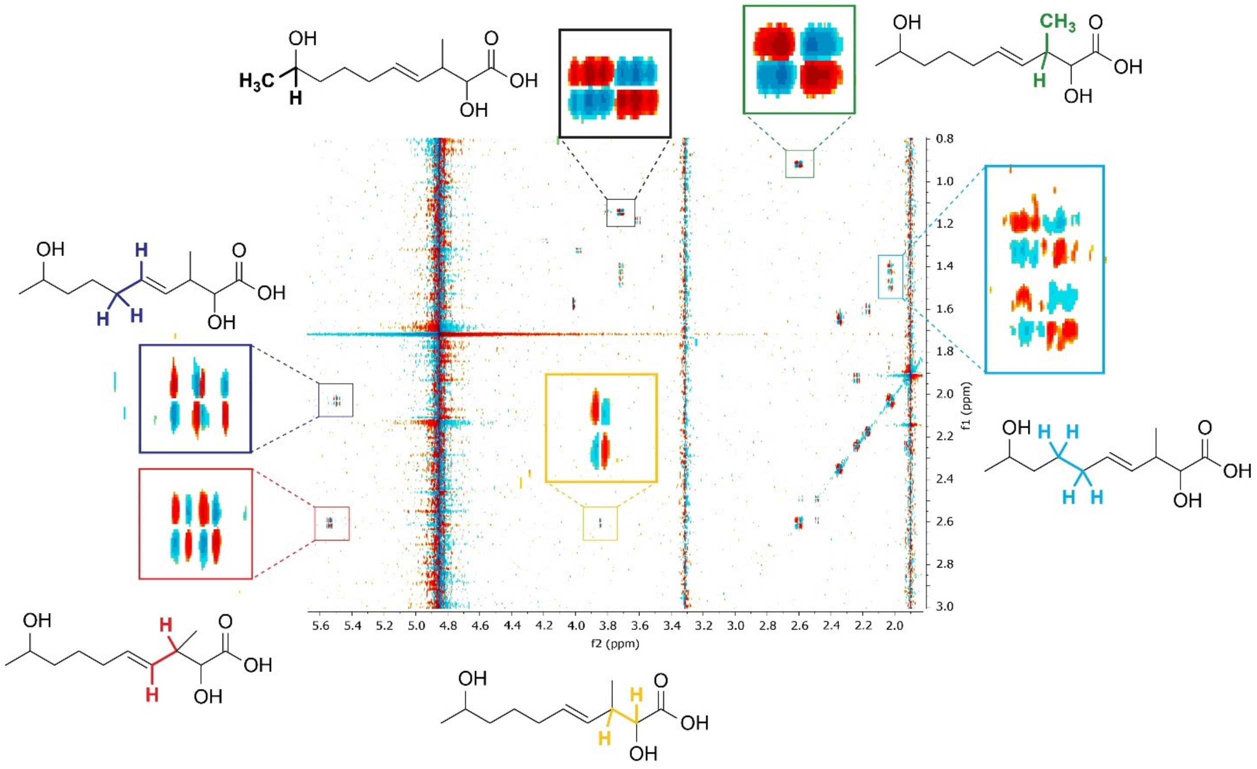

**Supplementary Figure 4.** Structure elucidation of bemeth#3.1. 2D NMR spectroscopic characterization of an isolated sample of compound **12** (bemeth#3.1, $C_{11}H_{20}O_4$) via dqfCOSY. Shown are relevant dqfCOSY cross peaks and the corresponding structural features.

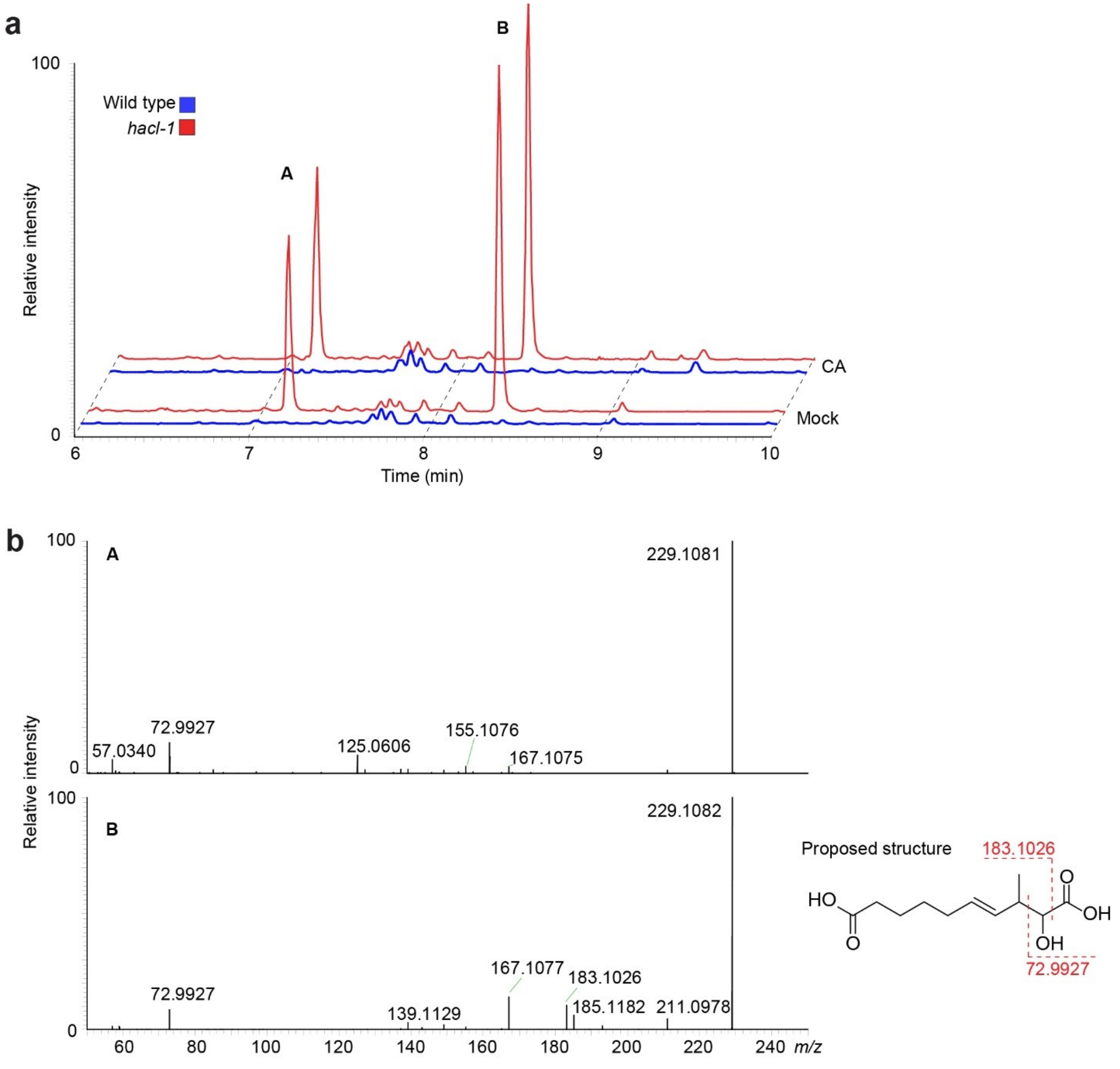

**Supplementary Figure 5.** Oxidized $C_{11}$ derivatives accumulate in *hacl-1*. **a**, Representative HPLC-MS (negative ion) EIC for *m/z* 229.1082, corresponding to $C_{11}H_{17}O_5^-$, from WT and *hacl-1* animals supplemented with CA or vehicle only, as indicated. Two major isomers are enriched in *hacl-1* irrespective of CA supplement, labeled A and B. **b**, MS/MS spectra for isomers of $C_{11}H_{17}O_5^-$, as indicated in panel (a). Features A and B both exhibit the diagnostic glyoxylate product ion (*m/z* 72.993), but only feature B also exhibits neutral loss of formic acid (*m/z* 183.1026). Proposed structure for B shown.

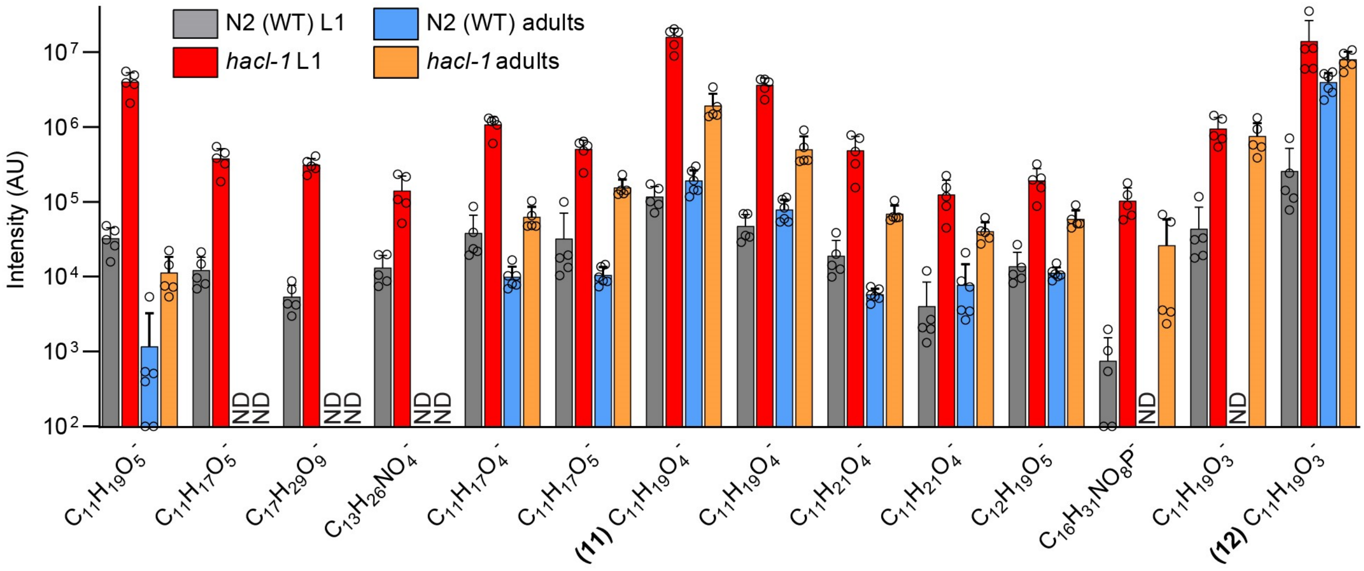

**Supplementary Figure 6**. Analysis of $C_{11}$ fatty acids in *C. elegans* larvae (L1) and adults. Quantification of metabolites in *exo*-metabolome extracts of synchronized N2 (WT) and *hacl-1* animals from the indicated stage. Quantified metabolites were originally identified as enriched in *hacl-1* L1 larvae *exo*-metabolome extracts (see **Figure 3**). Data for larvae represent five independent experiments and for adults represent six (N2) or five (*hacl-1*) samples from three biologically independent experiments and bars means ± standard deviation.

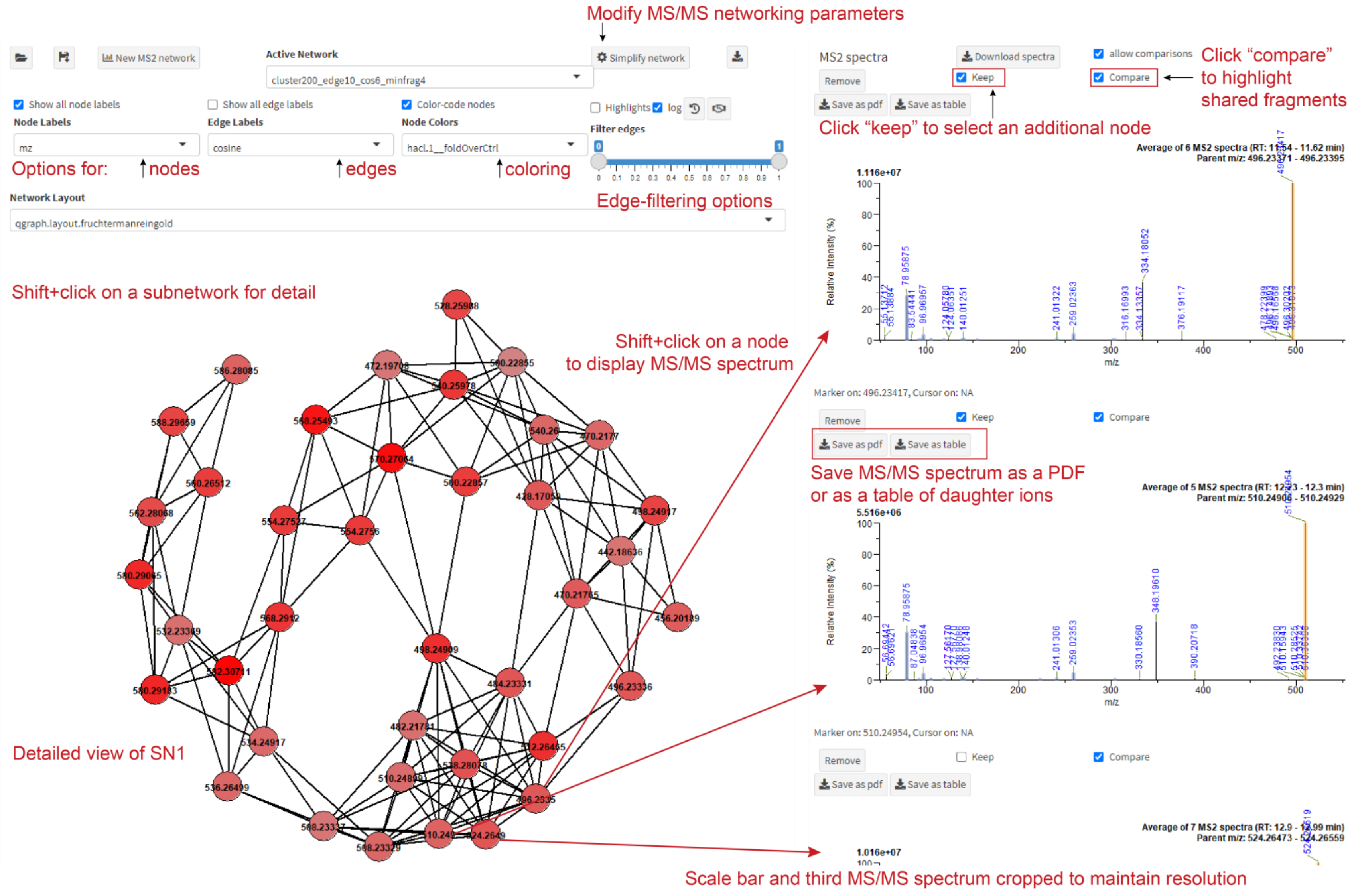

**Supplementary Figure 7.** Interact with MS/MS data using the *Keep and Compare* function. A screenshot of the *Compare MS2* module within the *Data Viewer*. Shift+click a subnetwork of interest for interactive capability, SN1 shown. MS/MS spectra displayed for *m/z* 496.2341, 510.2495, and 524.2652, corresponding to *N*-acyl GPE-13:1, -14:1, and -15:1, respectively. Toggling the *Keep* button allows users to display multiple MS/MS spectra by clicking additional nodes. Toggling the *Compare* button highlights the parental *m/z* in yellow and any shared fragments are highlighted in blue. MS/MS spectra are interactive: shift+click an *m/z* of interest to display all other *m/z* relative to selection. Neutral loss of a hexose moiety (-162.053) yields the major product ion in each spectrum. MS/MS spectra can be downloaded using the *Save As PDF* function; product ion tables can be downloaded in text format using the *Save As Table* function.

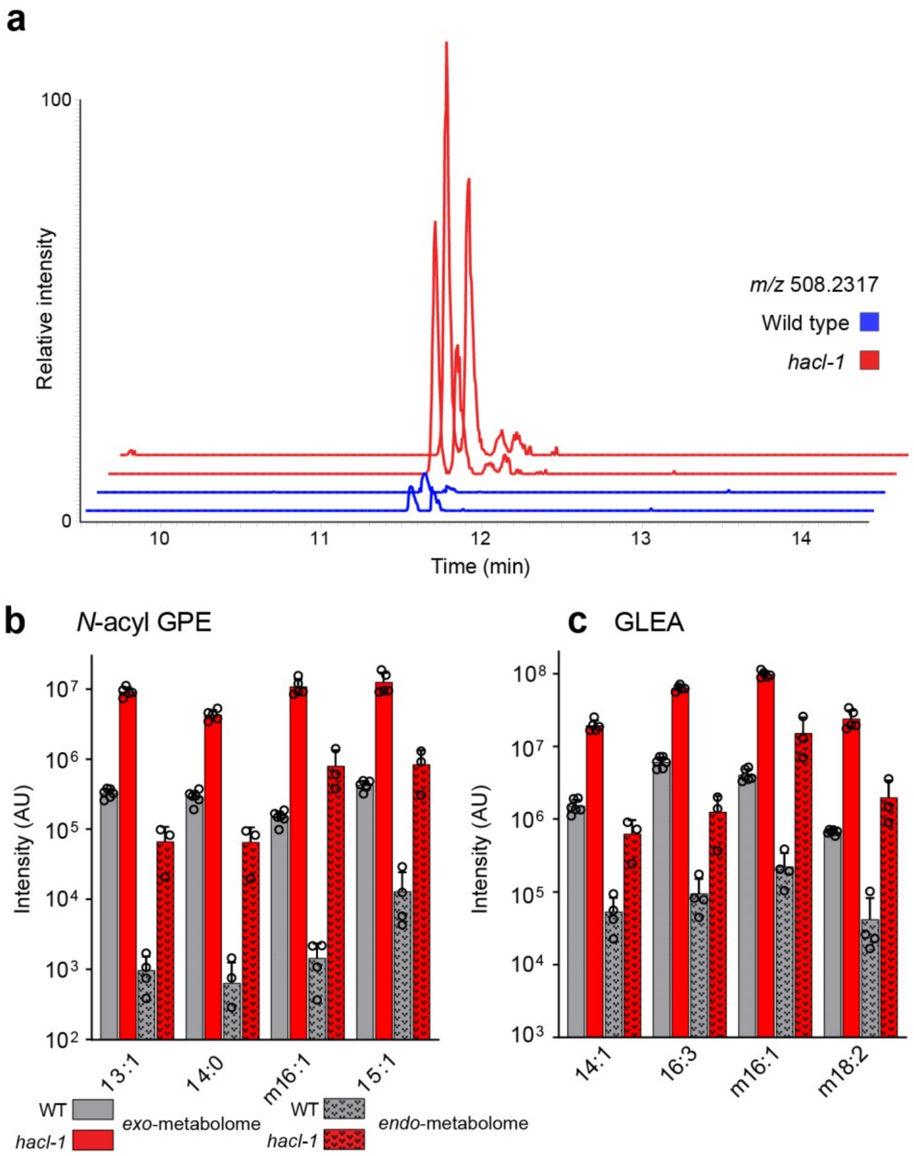

**Supplementary Figure 8.** *N*-acyl GPE and GLEA are more abundant in *exo*- than *endo*-metabolome. **a**, Representative HPLC-MS (negative ion) EIC for *m/z* 508.2317, corresponding to *N*-acyl GPE-14:2, in *exo*-metabolome extracts from N2 (WT) and *hacl-1*. Four distinct isomers of *N*-acyl GPE-14:2 observed (four peaks, two major, two minor). **b**, Quantification of representative *N*-acyl GPE from SN1 in the *exo*- and *endo*-metabolome extracts, as indicated. **c**, Quantification of representative GLEA from SN2 (14:1, 16:3) and SN3 (m16:1, m18:2) in the *exo*- and *endo*-metabolome extracts, as indicated. Data represent six (N2) or five (*hacl-1*) samples from three biologically independent experiments for *exo*-metabolome; data represent three biologically independent experiments for endo-metabolome and bars means ± standard deviation.

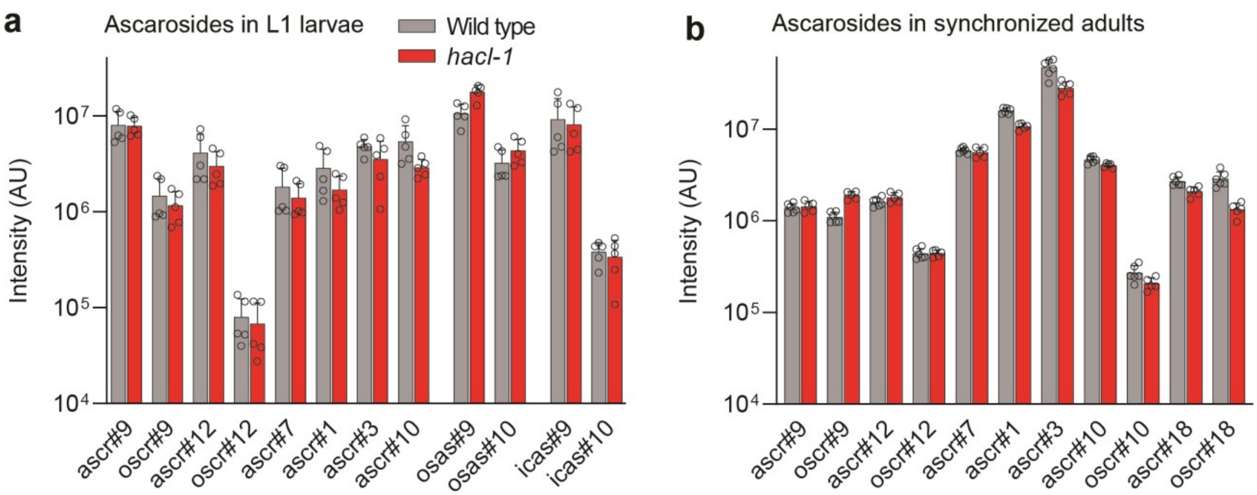

**Supplementary Figure 9.** Ascaroside biosynthesis is not significantly perturbed in *hacl-1* mutants. **a,** Quantification of representative ascarosides containing odd and even chain lengths from *exo*-metabolome extracts of N2 (WT) and *hacl-1* larvae, as indicated. Data represent five biologically independent experiments and bars means ± standard deviation. **b,** Quantification of representative ascarosides containing odd and even chain lengths from *exo*-metabolome extracts of N2 (WT) and *hacl-1* adults. Data represent six (N2) or five (*hacl-1*) samples from three biologically independent experiments and bars means ± standard deviation. Full structures of ascarosides listed can be accessed at the *C. elegans* Small Molecule Identifier Database (SMID-DB, www.smid-db.org). AU, arbitrary units

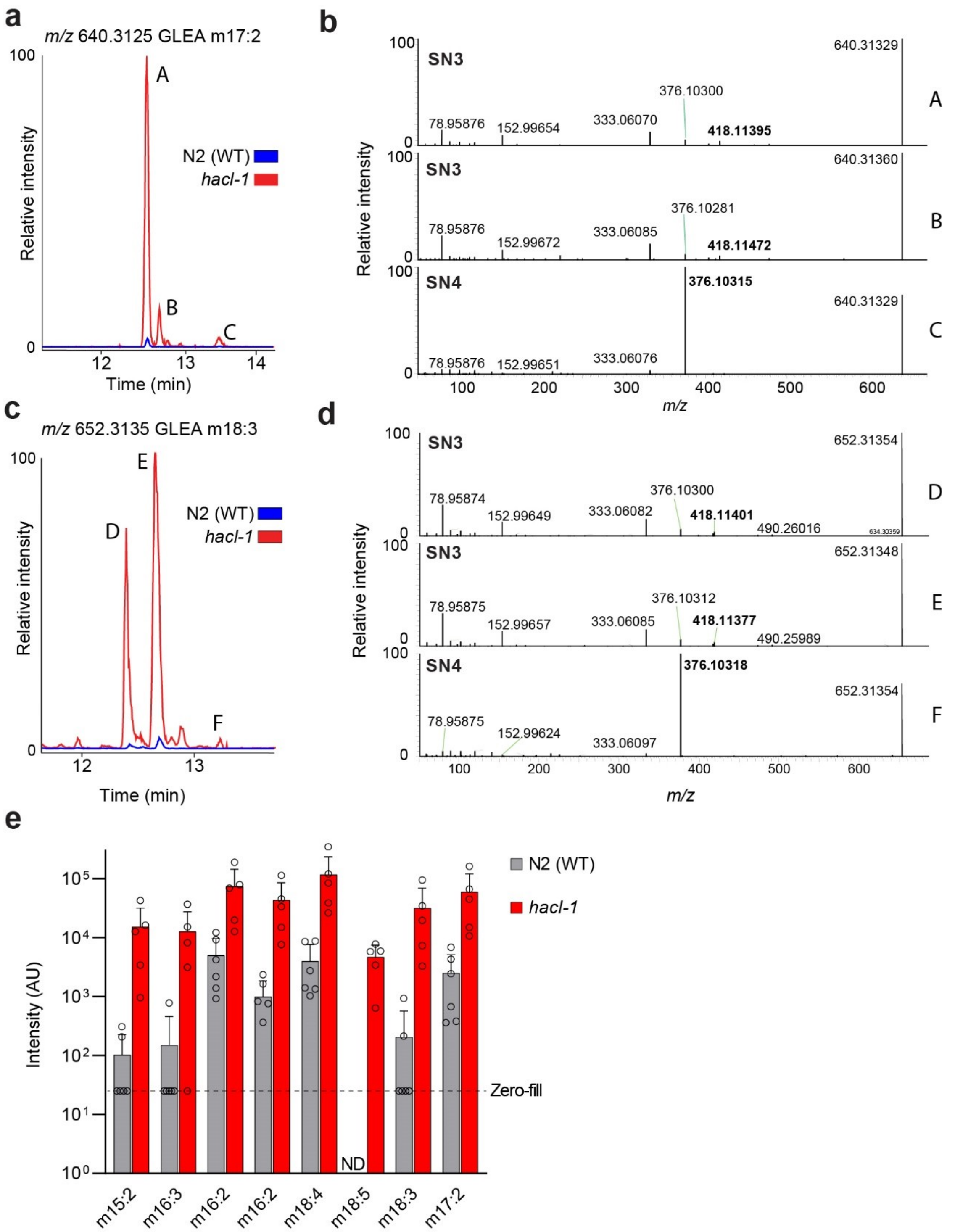

**Supplementary Figure 10.** GLEA in SN4 exhibit unique fragmentation and are low abundance. **a**, Representative HPLC-MS (negative ion) EIC for *m/z* 640.3125, corresponding to GLEA m17:2, which exhibits three major isomers, labeled as A, B, and C. **b,** MS/MS spectra for isomers of GLEA m17:2, as indicated in panel (a). Features A and B are networked in SN3 and produce the product ion *m/z* 418.114 during MS/MS fragmentation, whereas Feature C is networked in SN4 and produces an intense fragment ion *m/z* 376.103. **c,** Representative HPLC-MS (negative ion) EIC for *m/z* 652.3135, corresponding to GLEA m18:3, which exhibits five major isomers, three of which are labeled as D, E, and F. **d,** MS/MS spectra for isomers of GLEA m18:3, as indicated in panel (c). The major features D and E produce product ion *m/z* 418.114 during MS/MS fragmentation and are networked in SN3, whereas F belongs to SN4 and produces an intense fragment with *m/z* 376.103. **e**, Quantification of GLEA from SN4 in *exo*-metabolome extracts of N2 (WT) and *hacl-1*, as indicated. Data represent six (N2) or five (*hacl-1*) samples from three biologically independent experiments and bars means ± standard deviation. AU, arbitrary units, ND, not detected. The zero-fill line from value imputation is shown in cases where a feature was not detected in a subset of samples.

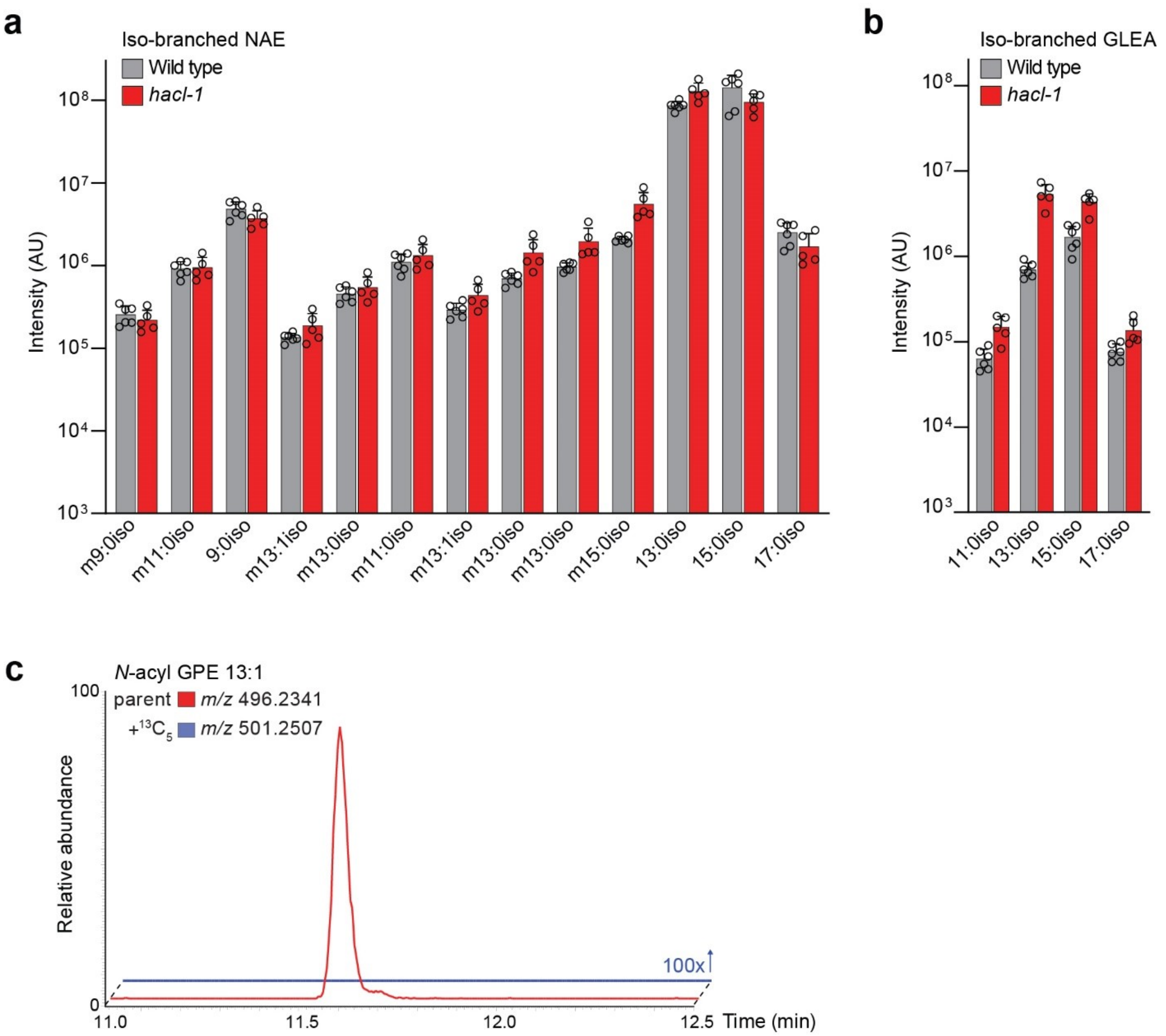

**Supplementary Figure 11.** Isotope tracing reveals BCFA-derived NAE and GLEA. Quantification of **a**, NAE and **b**, GLEA identified as $^{13}C_5$-enriched following $^{13}C_6$-Leu supplement from *exo*-metabolome extracts of N2 (WT) and *hacl-1* adults. Data represent six (N2) or five (*hacl-1*) samples from three biologically independent experiments and bars means ± standard deviation. **c**, Representative EICs for *m/z* 496.234 and 501.2507, corresponding to *N*-acyl GPE-13:1 and $^{13}C_5$- *N*-acyl GPE-13:1, from *exo*-metabolome extracts of N2 supplemented with $^{13}C_6$-Leu. Y-axis for *m/z* 501.2507 is scaled 100-fold to highlight absence of isotopic enrichment, demonstrating that this lipid is not a BCFA derived from Leu.

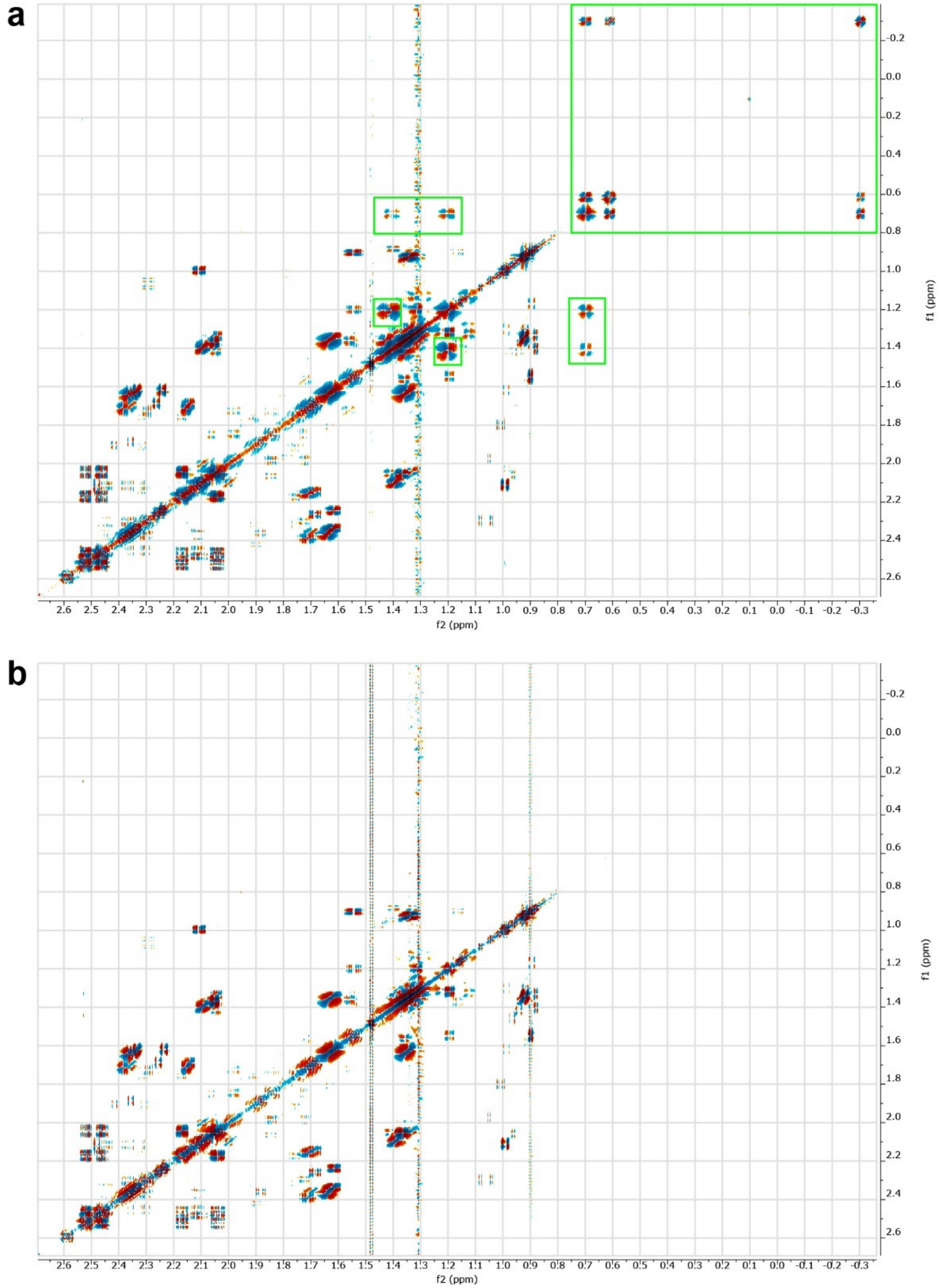

**Supplementary Figure 12.** JW1653-1 bacteria do not produce cyclopropane lipids. **a**, dqfCOSY spectrum (aliphatic region) of *C. elegans* reared on *E. coli* OP50 as food, showing crosspeaks characteristic for cyclopropane lipids (green boxes). **b,** dqfCOSY spectrum (aliphatic region) of *C. elegans* reared on JW1653-1 bacteria, lacking cyclopropyl signals. Spectra were acquired at 800 MHz, using $CD_3OD$ as solvent.

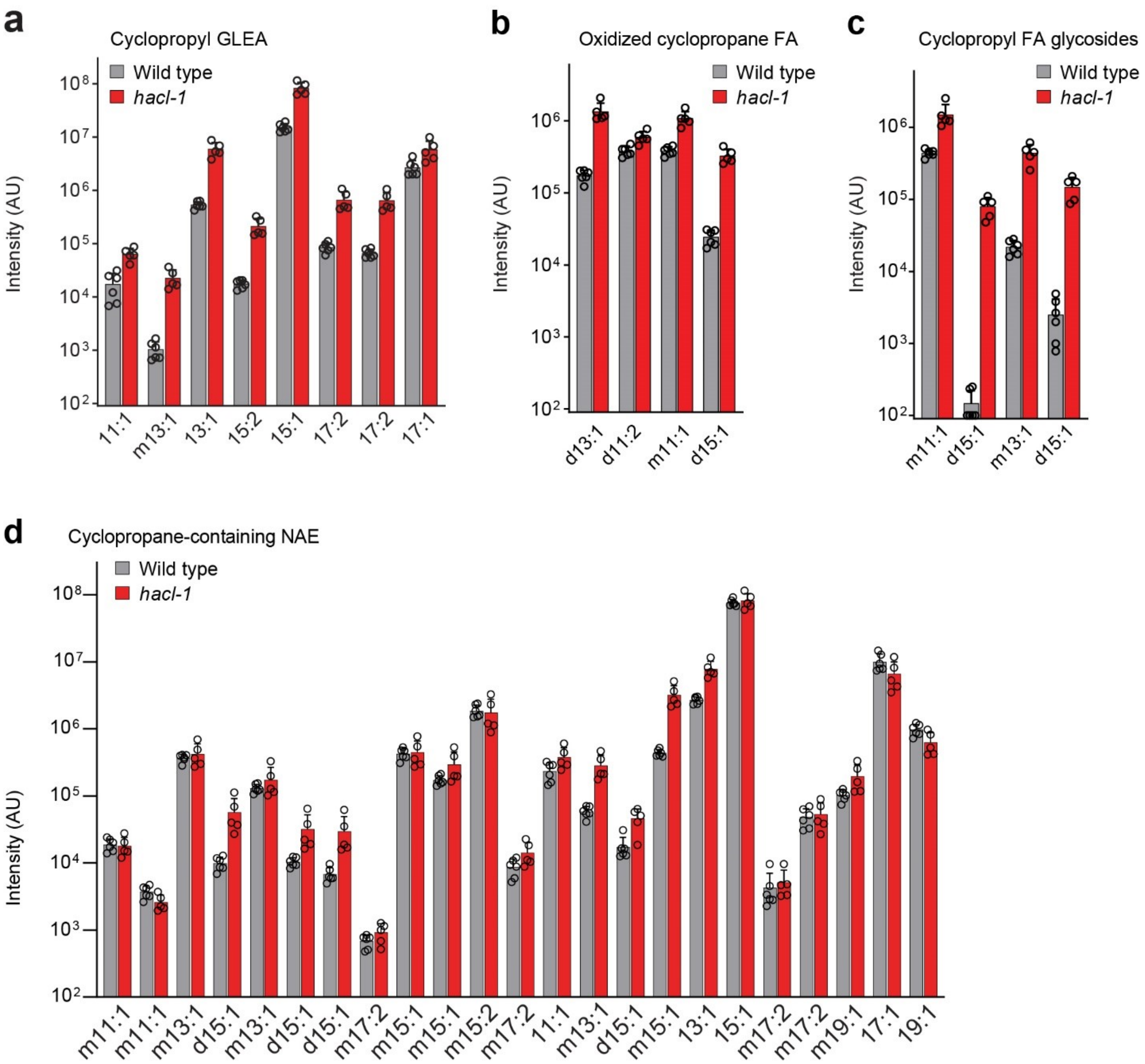

**Supplementary Figure 13.** Additional cyclopropane-containing metabolites identified by comparative analysis. Quantification of **a**, GLEA **b**, oxidized fatty acids **c**, fatty acyl glycosides and **d**, NAE that were absent from worms fed JW1653-1. Data represent six (N2) or five (*hacl-1*) samples from three biologically independent experiments and bars means ± standard deviation.

### Supplementary Tables

**Supplementary Table S1.** Comparison of Metaboseek to other free, open-source metabolomics tools with graphical user interfaces (GUI). Metaboseek offers flexibility in deployment as well as a unique combination of functionalities for discovery metabolomics.

| | Metaboseek | MZmine2 | XCMS online | MS-DIAL | GNPS Dashboard | MetaboAnalyst |
|---|---|---|---|---|---|---|
| **Deployment** | | | | | | |
| Server / local | Both | Local | Online | Local | Online | Online |
| Open-Source | + | + | Partial | + | + | Partial |
| Language | R/shiny | Java | R/Java script | C# | Python | R/PrimeFaces |
| **Raw Data Processing** | | | | | | |
| Feature detection | + | + | + | + | Limited | + |
| Interactive raw data browser | + | + | - | + | + | - |
| Grouped EIC plot | + | - | Limited | + | - | - |
| **Feature Tables** | | | | | | |
| Import feature tables | + | + | - | - | - | + |
| Statistical analysis | + | + | + | + | - | + |
| Custom sample group filters | + | - | - | - | - | - |
| **Structural Characterization** | | | | | | |
| Formula prediction / database lookup | + | + | + | + | + | + |
| Molecular Networking | + | Export | Export | + | + | - |
| SIRIUS fragment annotation | + | + | - | - | - | - |
| MS2 pattern search | + | + | - | + | - | - |
| Isotope Tracking | + | - | - | + | - | - |

**Supplementary Table S2. NMR spectroscopic data of bemeth#3 (11)** (600 MHz, methanol-$d_4$).

| Position | δ $^1$H [ppm] | $^1$H-$^1$H-coupling constants [Hz] |
|---|---|---|
| 1 | | |
| 2 | 3.84 | $J_{2,3}$ = 3.8 Hz |
| 3 | 2.60 | $J_{2,3}$ = 8.5 Hz |
| 4 | 5.53 | $J_{4,5}$ = 16 Hz |
| 5 | 5.50 | $J_{5,6}$ = 7.3 Hz |
| 6 | 2.02 | |
| 7 | 1.40 | |
| | 1.48 | |
| 8 | 1.41 | |
| | 1.47 | |
| 9 | 3.72 | $J_{9,10}$ = 6.3 |
| 10 | 1.17 | |
| 3-$CH_3$ | 0.92 | $J_{3,3\text{-}CH3}$ = 6.9 |

**Supplementary Table S3. NMR spectroscopic data of bemeth#2 (12), major diastereomer** (600 MHz, methanol-$d_4$).

| Position | δ $^{13}$C [ppm] | δ $^1$H [ppm] | $^1$H-$^1$H-coupling constants [Hz] | HMBC correlations |
|---|---|---|---|---|
| 1 | 179.3 | | | |
| 2 | 76.8 | 3.86 | $J_{2,3}$ = 3.8 Hz | C-1 (weak), C-11 |
| 3 | 41.5 | 2.59 | | |
| 4 | 134.4 | 5.50 | | C-3, C-6, C-11 |
| 5 | 130.7 | 5.49 | | C-3, C-6 |
| 6 | 33.4 | 2.00 | $J_{5,6}$ = 7.3, $J_{6,7}$ = 6.0 | C-4, C-5, C-7, C-8 |
| 7 | 30.1 | 1.38 | | C-5, C-6, C-8, C-9 |
| 8 | 32.3 | 1.31 | | C-6, C-7, C-9, C-10 |
| 9 | 23.4 | 1.32 | $J_{9,10}$ = 7.1 | C-7, C-8, C-10 |
| 10 | 14.1 | 0.90 | | C-8, C-9 |
| 3-$CH_3$ | 14.2 | 0.92 | $J_{3,3\text{-}CH3}$ = 6.9 | C-2, C-3, C-4 |

**Supplementary Table S4.** NMR spectroscopic data for GLEA-m16:1 (**16**), methanol-$d_4$ (800 MHz).

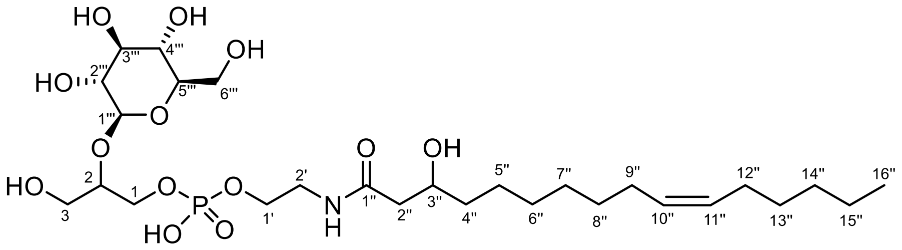

**16** Proposed structure, position of double bond not known

| Position | δ $^{13}$C [ppm] | δ $^{1}$H [ppm] ($J_{HH}$/$J_{HP}$ [Hz]) | Key HMBC signals |
|---|---|---|---|
| 1’’ | 174.2 | | |
| 2’’a | 44.5 | 2.31 (dd, $J_{2a'',2b''}$ = 15.0, $J_{2a'',3''}$ = 8.1) | C-1’’ |
| 2’’b | | 2.36 (dd, $J_{2b'',3''}$ = 4.4) | C-1’’ |
| 3’’ | 69.4 | 3.96 (m) | |
| 4’’ | 38.0 | 1.46 (m, 2 H) | |
| 5’’-8’’ | 29.8-30.7 (4 C) | 1.32-1.38 (m, 8 H) | |
| 9’’/12’’ | 27.8 | 2.02-2.06 (m, 4 H) | |
| 10’’/11’’ | 130.5 (2C) | 5.33-5.36 (m, 2H) | |
| 13’’-15’’ | 29.8-30.7 (3 C) | 1.29-1.38 (m, 6 H) | |
| 16’’ | 14.1 | 0.90 (t, $J_{15'',16''}$ = 7.0) | |
| 1′ | 64.7 | 3.93 (dt, $J_{1',P}$ = 6.5, $J_{1',2'}$ = 5.8) | |
| 2′ | 41.1 | 3.42 (t, $J_{1',2'}$ = 5.8) | C-1’’ |
| 1a | 65.4 | 3.93 (ddd, $J_{1a,2}$ = 6.8, $J_{1a,1b}$ = 10.7, $J_{1a,P}$ = 6.3) | |
| 1b | | 4.06 (ddd, $J_{1b,2}$ = 3.8, $J_{1b,P}$ = 6.3) | |
| 2 | 80.5 | 3.90 (m) | C-1’’’ (weak) |
| 3a | 62.2 | 3.69 (dd, $J_{2,3a}$ = 6.0, $J_{3a,3b}$ = 12.5) | |
| 3b | | 3.78 (dd, $J_{2,3b}$ = 3.4) | |
| 1′′′ | 104.2 | 4.43 (d, $J_{1''',2'''}$ = 8) | C-2 |
| 2′′′ | 75.0 | 3.22 (dd, $J_{3''',4'''}$ = 9) | C-1’’’ |
| 3′′′ | 77.6 | 3.36 (m) | |
| 4′′′ | 71.3 or 77.9 | 3.26-3.29 (m) | |
| 5′′′ | 71.3 or 77.9 | 3.26-3.29 (m) | |
| 6a′′′ | 62.4 | 3.67 (dd, $J_{5''',6a'''}$ = 5.6, $J_{6a''',6b'''}$ = 12) | |
| 6b′′′ | | 3.85 (dd, $J_{5''',6b'''}$ = 2) | |

### NMR Appendix

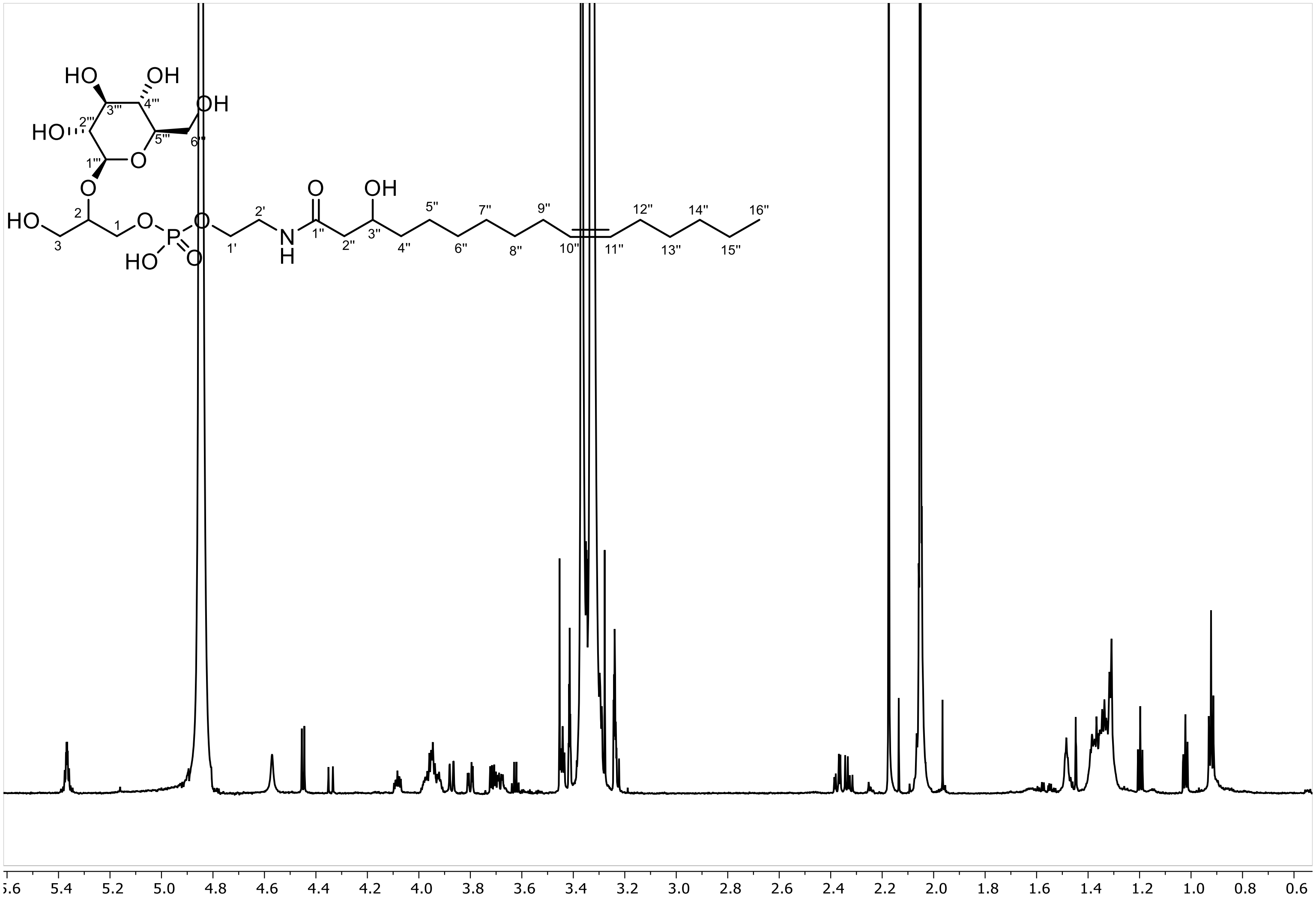

$^{1}$H NMR spectrum of isolated fraction containing GLEA-m16:1 (**16**), methanol-$d_4$ (800 MHz)

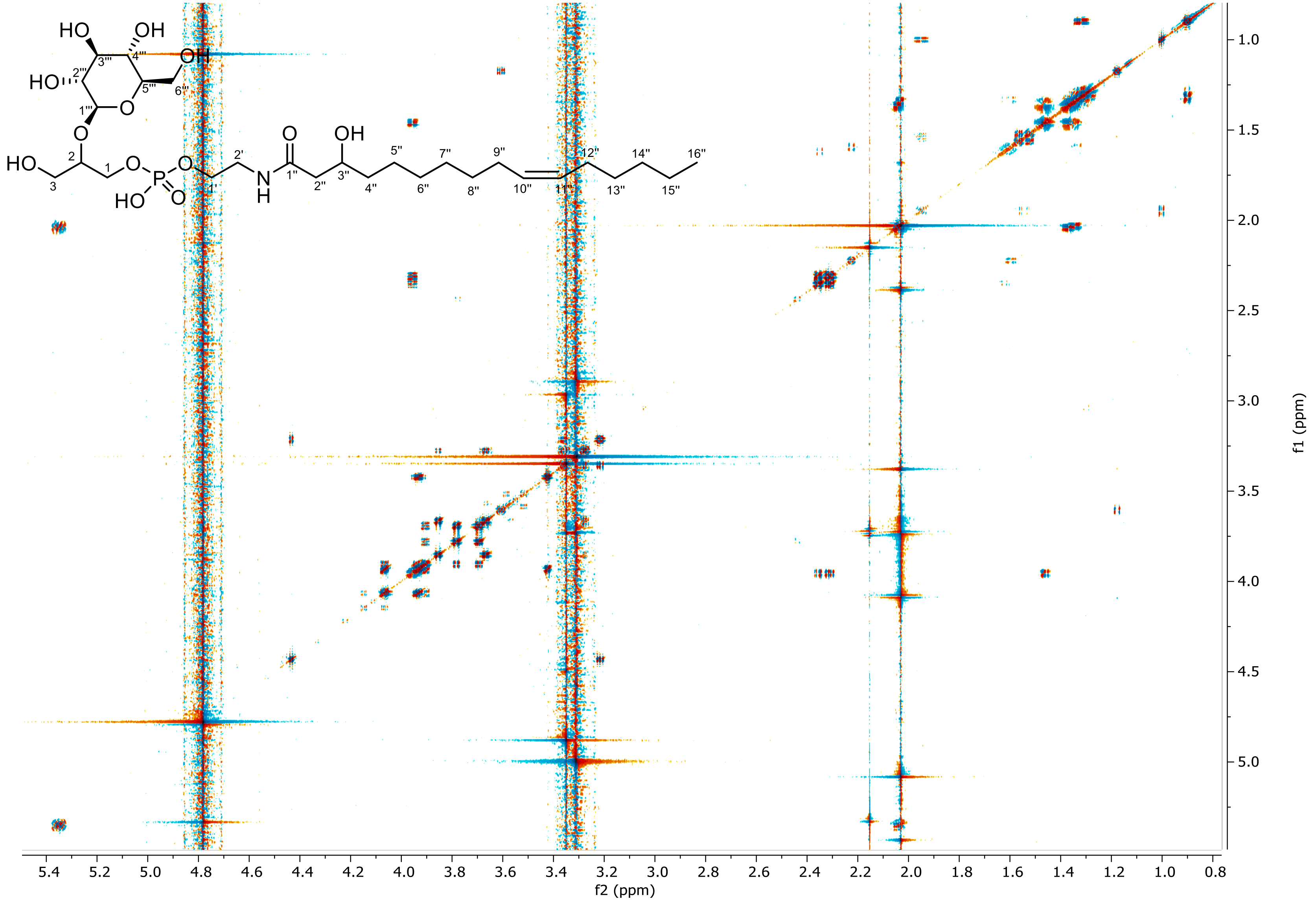

dqfCOSY spectrum of metabolome fraction containing GLEA-m16:1 (**16**), methanol-$d_4$ (800 MHz)

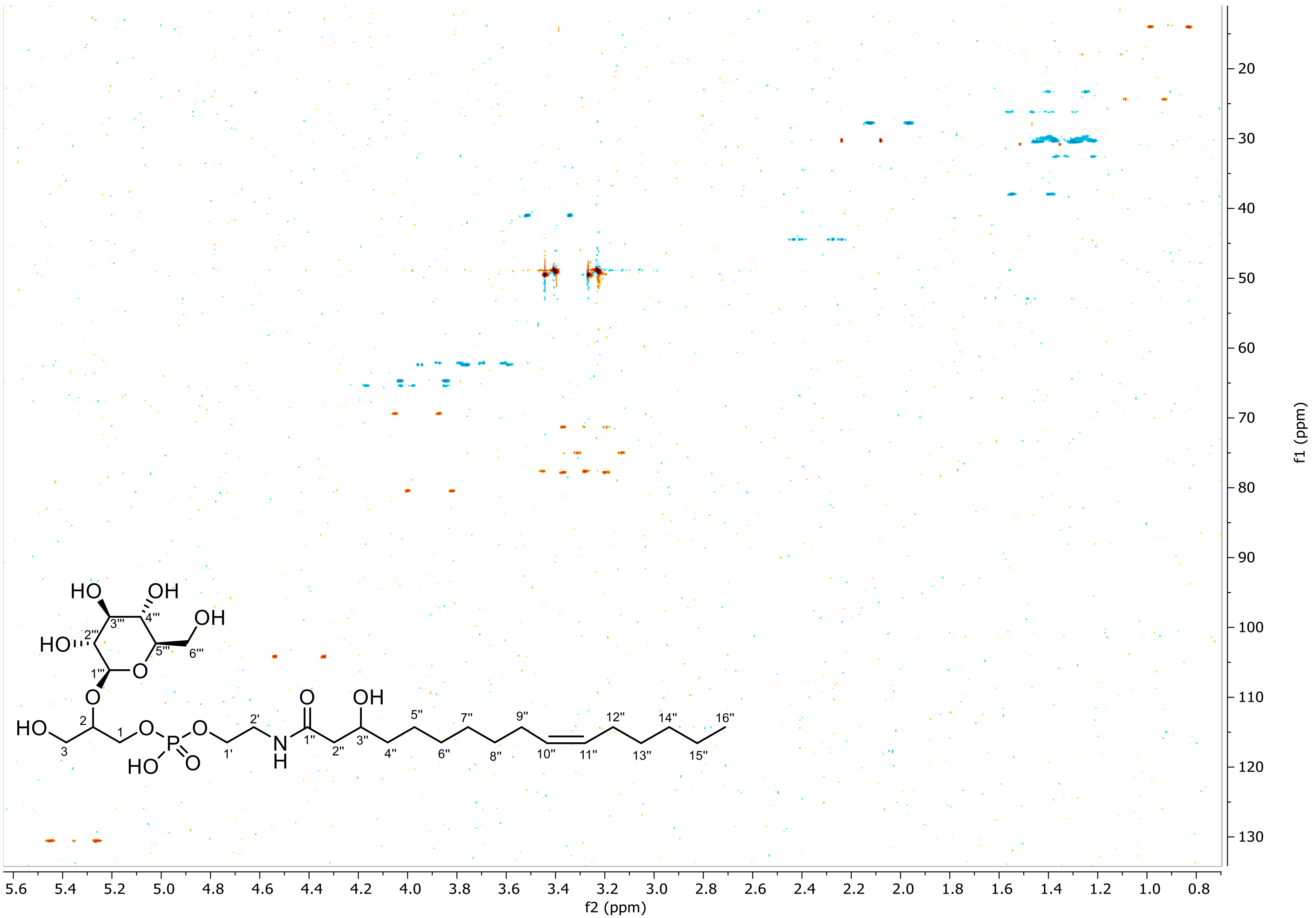

HSQC spectrum of metabolome fraction containing GLEA-m16:1 (**16**), methanol-$d_4$ (800 MHz)

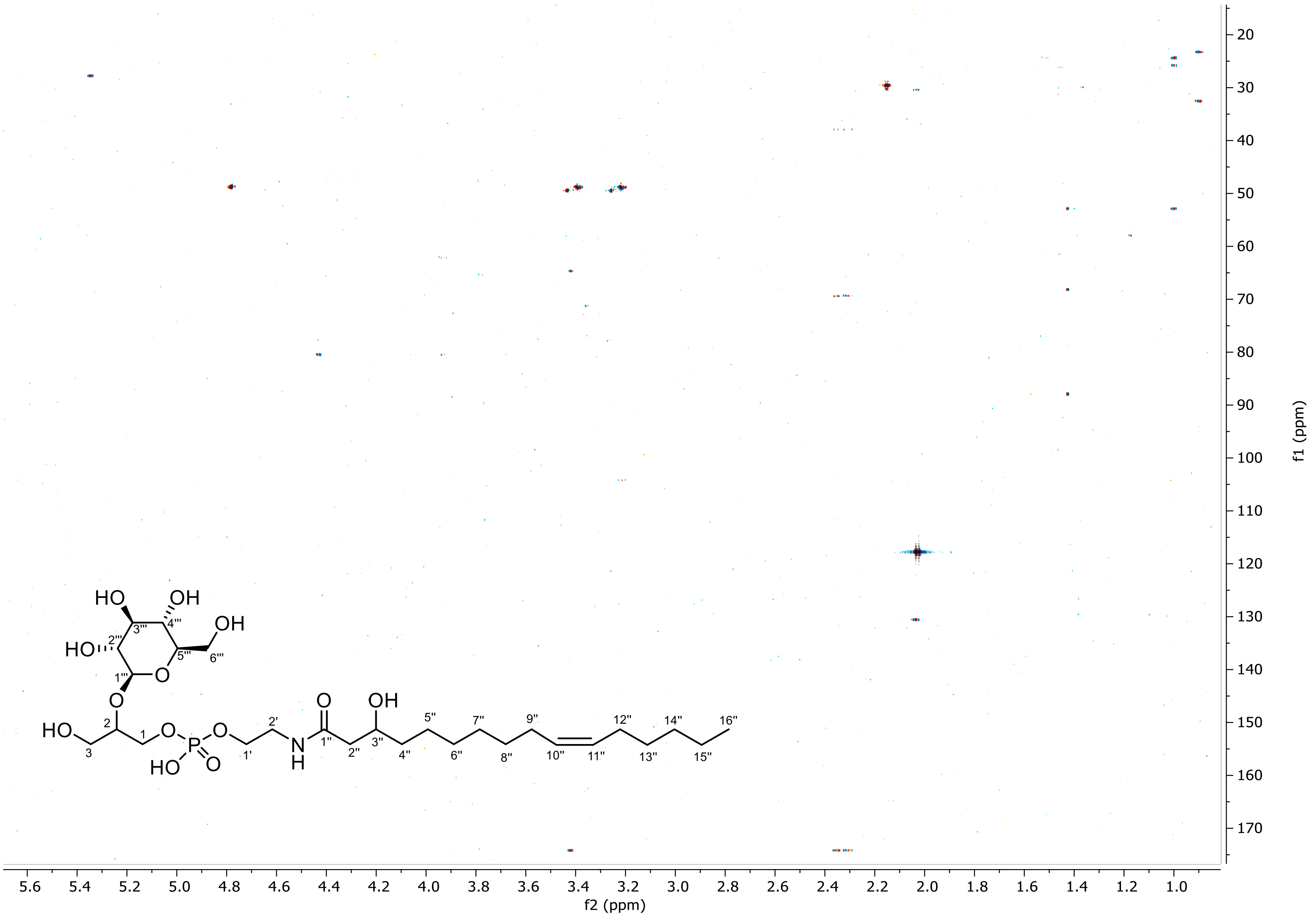

HMBC spectrum of metabolome fraction containing GLEA-m16:1 (**16**), methanol-$d_4$ (800 MHz)
